# Gene expression imputation across multiple brain regions reveals schizophrenia risk throughout development

**DOI:** 10.1101/222596

**Authors:** Laura M. Huckins, Amanda Dobbyn, Douglas M. Ruderfer, Gabriel Hoffman, Weiqing Wang, Antonio Pardinas, Veera M Rajagopal, Thomas D. Als, Hoang Nguyen, Kiran Girdhar, James Boocock, Panos Roussos, Menachem Fromer, Robin Kramer, Enrico Domencini, Eric Gamazon, Shaun Purcell, CommonMind Consortium, the Schizophrenia Working Group of the Psychiatric Genomics Consortium, iPSYCH-GEMS Schizophrenia Working Group, Ditte Demontis, Anders D. Børglum, James Walters, Michael O’Donovan, Patrick Sullivan, Michael Owen, Bernie Devlin, Solveig K. Sieberts, Nancy Cox, Hae Kyung Im, Pamela Sklar, Eli A. Stahl

## Abstract

Transcriptomic imputation approaches offer an opportunity to test associations between disease and gene expression in otherwise inaccessible tissues, such as brain, by combining eQTL reference panels with large-scale genotype data. These genic associations could elucidate signals in complex GWAS loci and may disentangle the role of different tissues in disease development. Here, we use the largest eQTL reference panel for the dorso-lateral pre-frontal cortex (DLPFC), collected by the CommonMind Consortium, to create a set of gene expression predictors and demonstrate their utility. We applied these predictors to 40,299 schizophrenia cases and 65,264 matched controls, constituting the largest transcriptomic imputation study of schizophrenia to date. We also computed predicted gene expression levels for 12 additional brain regions, using publicly available predictor models from GTEx. We identified 413 genic associations across 13 brain regions. Stepwise conditioning across the genes and tissues identified 71 associated genes (67 outside the MHC), with the majority of associations found in the DLPFC, and of which 14/67 genes did not fall within previously genome-wide significant loci. We identified 36 significantly enriched pathways, including hexosaminidase-A deficiency, and multiple pathways associated with porphyric disorders. We investigated developmental expression patterns for all 67 non-MHC associated genes using BRAINSPAN, and identified groups of genes expressed specifically pre-natally or post-natally.

## Introduction

Genome-wide association studies (GWAS) have yielded large lists of disease-associated loci. Despite this, progress in identifying the causal variants driving these associations, particularly for complex psychiatric disorders such as schizophrenia, has lagged much further behind. Interpreting associated variants and loci is therefore vital to understanding how genetic variation contributes to disease pathology. Expression Quantitative Trait Loci (eQTLs), which are responsible for a substantial proportion of gene expression variance, have been posited as a potential link between associated loci and disease susceptibility^1–5^, and indeed have yielded results for a host of complex traits^6–9^. Consequently, numerous methods to identify and interpret co-localisation of eQTLs and GWAS loci have been developed^10–13^. However, these methods require simplifying assumptions about genetic architecture (i.e., one causal variant per GWAS locus) and/or linkage disequilibrium, may be underpowered or overly conservative, especially in the presence of allelic heterogeneity, and have not yet yielded substantial insights into existing or novel loci.

Biologically relevant information can be extracted by transcriptomic investigations, as recently described by the CommonMind Consortium^14^ (CMC), thanks to detailed RNA-sequencing in a large cohort of genotyped individuals with schizophrenia and bipolar disorder^14^. These analyses however are underpowered to detect with statistical confidence differential expression of genes mapping at schizophrenia (SCZ) risk loci, due to the small effects predicted by GWAS combined with the difficulty of obtaining adequate sample sizes of neurological tissues^14^. Still, such methods do not necessarily identify all risk variation in GWAS loci. Transcriptomic imputation is an alternative approach that leverages large eQTL reference panels to bridge the gap between large-scale genotyping studies and biologically useful transcriptome studies^15,16^. This approach seeks to identify and codify the relationships between genotype and gene expression in matched panels of individuals, then impute the genetic component of the transcriptome into large-scale genotype-only datasets, such as case-control GWAS cohorts, which enables investigation of disease-associated gene expression changes. This will allow us to study genes with modest effect sizes, likely representing a large proportion of genomic risk for psychiatric disorders^14,17^.

The access to the large collection of dorso-lateral pre-frontal cortex (DLPFC) gene expression data collected by the CommonMind Consortium^14^ affords us a unique opportunity to study and codify relationships between genotype and gene expression. Here, we present a novel set of gene expression predictor models, built using CommonMind Consortium DLPFC data^14^. We compare different regression approaches to building these models (including elastic net^15^, Bayesian sparse linear mixed models and ridge regression^16^, and using max eQTLs), and benchmark performance of these predictors against existing GTEx prediction models. We applied our CMC DLPFC predictors and 12 GTEx-derived neurological prediction models to predict gene expression in schizophrenia GWAS data, obtained through collaboration with the Psychiatric Genomics Consortium (PGC) schizophrenia working group, the “CLOZUK2” cohort, and the iPSYCH-GEMS schizophrenia working group. We identified 413 genome-wide significant genic associations with schizophrenia in our PGC+CLOZUK2 sample, constituting 67 independent associations outside the MHC region. We demonstrate the relevance of these associations to schizophrenia aetiopathology using gene set enrichment analysis, and by examining the effects of manipulation of these genes in mouse models. Finally, we investigated spatio-temporal expression of these genes using a developmental transcriptome dataset, and identified distinct spatio-temporal patterns of expression across our associated genes.

## Results

### Prediction Models based on CommonMind Consortium DLPFC expression

Using matched genotype and gene expression data from the CommonMind Consortium Project, we developed DLPFC genetically regulated gene expression (GREX) prediction models. We systematically compared four approaches to building predictors^15,16^ within a cross-validation framework. Elastic net regression had a higher distribution of cross-validation R^2^ (R_CV_^2^) and higher mean R_CV_^2^ values (Supplementary Figures 1, 2A) than all other methods. We therefore used elastic net regression to build our prediction models. We compared prediction models created using elastic net regression on SVA-corrected and uncorrected data^14^. The distribution of R_CV_^2^ values for the SVA-based models was significantly higher than for the un-corrected data^14,18^ (ks-test; p<2.2e-16; Supplementary figure 1B-C). In total, 10,929 genes were predicted with elastic net cross-validation R_CV_^2^ > 0.01 in the SVA-corrected data and were included in the final predictor database (mean R_CV_^2^ = 0.076).

**Figure 1:**
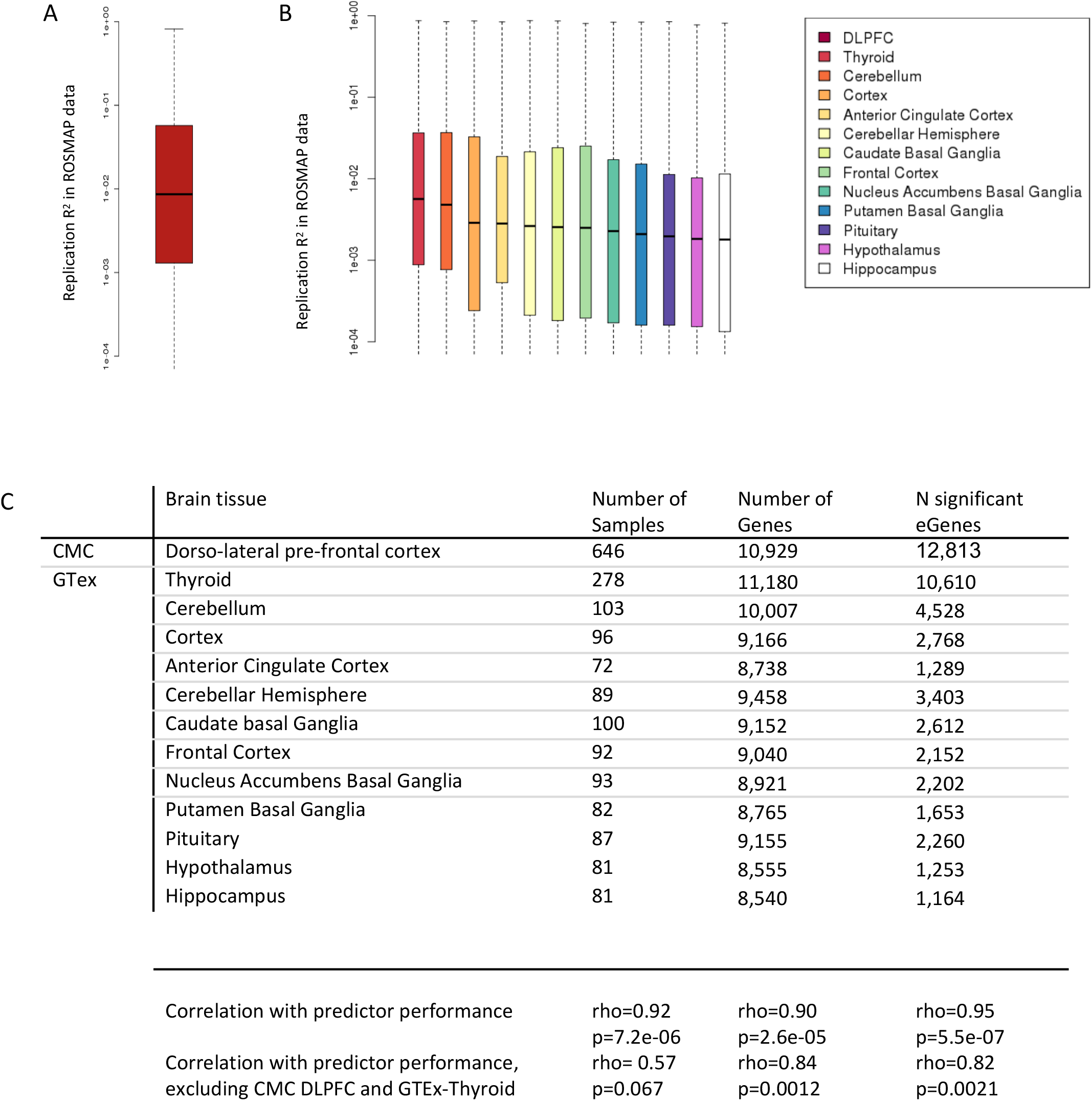
Replication of DLPFC prediction models in independent data. Measured gene expression (ROSMAP RNA-seq) was compared to predicted genetically-regulated gene expression for CMC DLPFC and 12 GTeX predictor databases. Replication R^2^ values are significantly higher for the DLPFC than for the 12 GTEX brain expression models. A. Distribution of Revalues of CMC DLPFC predictors in ROSMAP data. Mean R_R_^2^ = 0.056. 47.7% of genes have R_R_^2^ >= 0.01. B. Distribution of R_R_^2^ values of 12 GTeX predictors in ROSMAP data. C. Table of sample sizes and p-val thresholds for CMC DLPFC and GTeX data. Number of samples, number of genes in the prediXcan model and number of eGenes are all significantly correlated with predictor performance in ROSMAP data.

**Figure 2:**
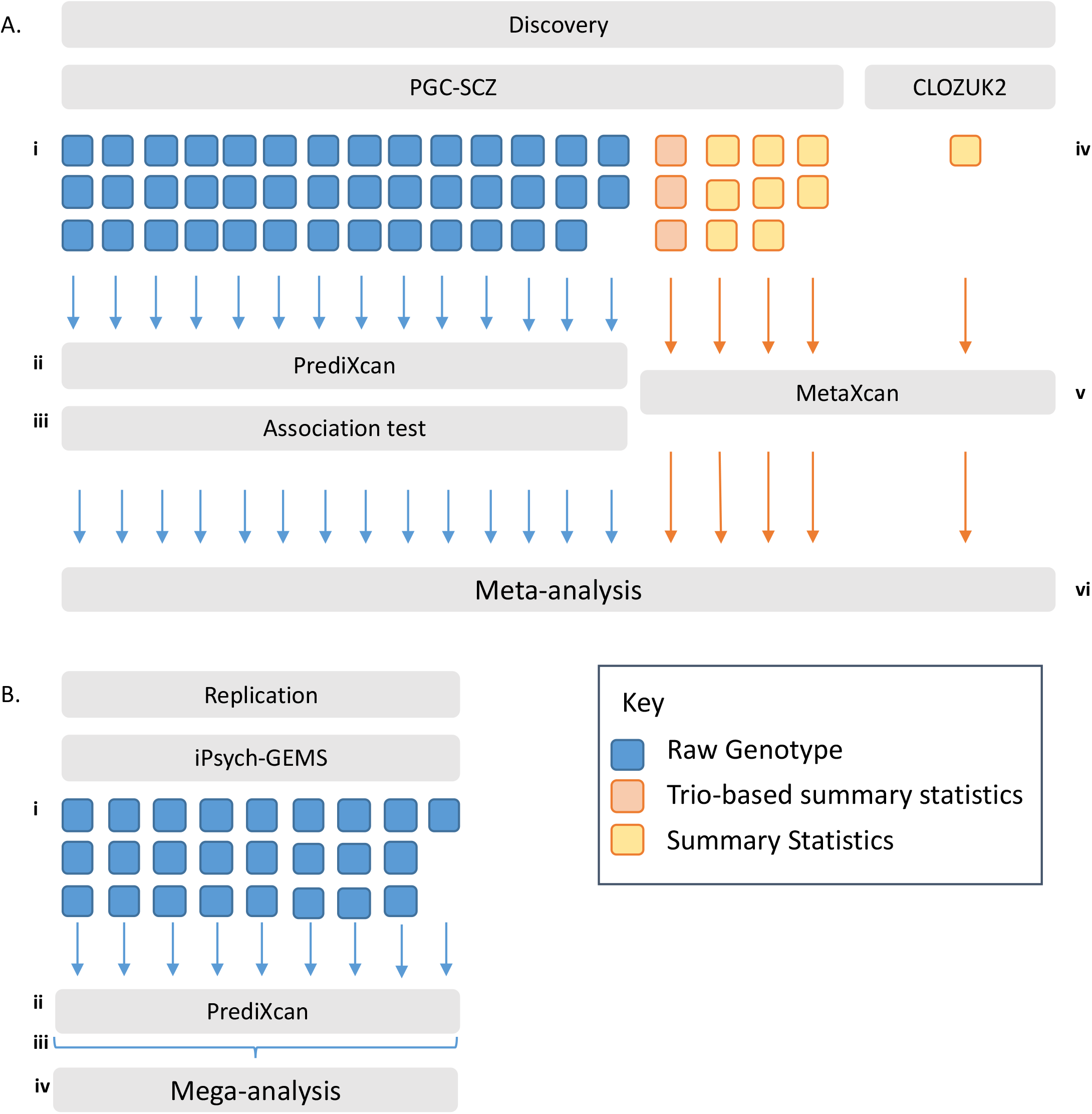
Analysis outline. A) Discovery Samples. 41 PGC-SCZ cohorts had available raw genotypes (i). Predicted DLPFC gene expression was calculated in each cohort using prediXcan (ii) and tested for association with case-control status (iii). 11 PGC cohorts (3 trio, 8 case-control) and the CLOZUK2 cohort had only summary statistics available (iv). MetaXcan was used to calculate DLPFC associations for each cohort (v). Results were meta-analysed across all 53 cohorts (vi). This procedure was repeated for 12 GTEx prediction models. B) Replication Samples. iPSYCH-GEMS samples were collected in 25 waves (i). Predicted DLPFC gene expression was calculated in each wave separately using prediXcan (ii) and merged for association testing (iii). A mega-analysis was run across all 25 waves, using wave membership as a covariate in the regression (iv)

To test the predictive accuracy of the CMC-derived DLPFC models, and to benchmark this against existing GTEx-derived prediction models, genetically-regulated gene expression (GREX) was calculated in an independent DLPFC RNA-sequencing dataset (the Religious Orders Study Memory and Ageing Project, ROSMAP^19^). We compared predicted GREX to measured ROSMAP gene expression for each gene (Replication R^2^, or R_R_^2^) for the CMC-derived DLPFC models and twelve GTEx-derived brain tissue models^15,20,21^ (Figure 1, Supplementary Figure 2B). CMC-derived DLPFC models had higher average R_R_^2^ values (Mean R_R_^2^ = 0.056), more genes with R_R_^2^ > 0.01, and significantly higher overall distributions of R_R_^2^ values than any of the twelve GTEx models (ks-test, p<2.2×10^−16^ across all analyses; Figure 1). Median R_R_^2^ values were significantly correlated with sample size of the original tissue set (rho=0.92, p=7.2×10^−6^), the number of genes in the prediction model (rho=0.9, p=2.6×10^−5^), and the number of significant ‘eGenes’ in each tissue type (rho=0.95, p=5.^5x10-7^; Figure 1C). Notably, these correlations persist after removing obvious outliers (Figure 1C).

To estimate trans-ancestral prediction accuracy, genetically regulated gene expression was calculated for 162 African-American individuals and 280 European individuals from the NIMH Human Brain Collection Core (HBCC) dataset (supplementary figure 2B). R_R_^2^ values were higher on average in Europeans than African-Americans (average R_R_EUR_^2^ = 0.048, R_R_AA_^2^ = 0.040), but were significantly correlated between African-Americans and Europeans (rho=0.78, p<2.2 ×10^−16^, Pearson test; supplementary figure 3).

### Application of Transcriptomic Imputation to Schizophrenia

We used CMC DLPFC and the 12 GTEx-derived brain tissue prediction models to impute genetically regulated expression levels (GREX) of 19,661 unique genes in cases and controls from the PGC-SCZ GWAS study^22^. Predicted expression levels were tested for association with schizophrenia. Additionally, we applied CMC and GTEx-derived prediction models to summary statistics from 11 PGC cohorts (for which raw genotypes were unavailable) and the CLOZUK2 cohort. Meta-analysis was carried out across all PGC-SCZ and CLOZUK2 cohorts using an odds-ratio based approach in METAL. Our final analysis included 40,299 cases and 65,264 controls (Figure 2A).

We identified 413 genome-wide significant associations, representing 256 genes in 13 tissues (Figure 3A). The largest number of associations were detected in the CMC DLPFC GREX data (Figure 3C; 49 genes outside the MHC, 69 genes overall). We sought replication of our CMC DLPFC SCZ-associations in an independent dataset of 4,133 cases and 24,788 controls in collaboration with the iPSYCH-GEMS SCZ working group (Figure 2B). We found significant correlation of effect sizes (p=1.784 ×10^−04^; rho=0.036) and −log10 p-values (p=1.073 ×10^−05^; rho=0.043) between our discovery (PGC+CLOZUK2) and replication (iPSYCH-GEMS) samples. Non-MHC Genes reaching genome-wide significance in our discovery sample (49 genes) were significantly more likely to reach nominal significance in the replication sample, and had significantly more consistent directions of effect than might be expected by chance (binomial test, p=2.42 ×10^−05^, p=0.044). (Suppl. info).

**Figure 3:**
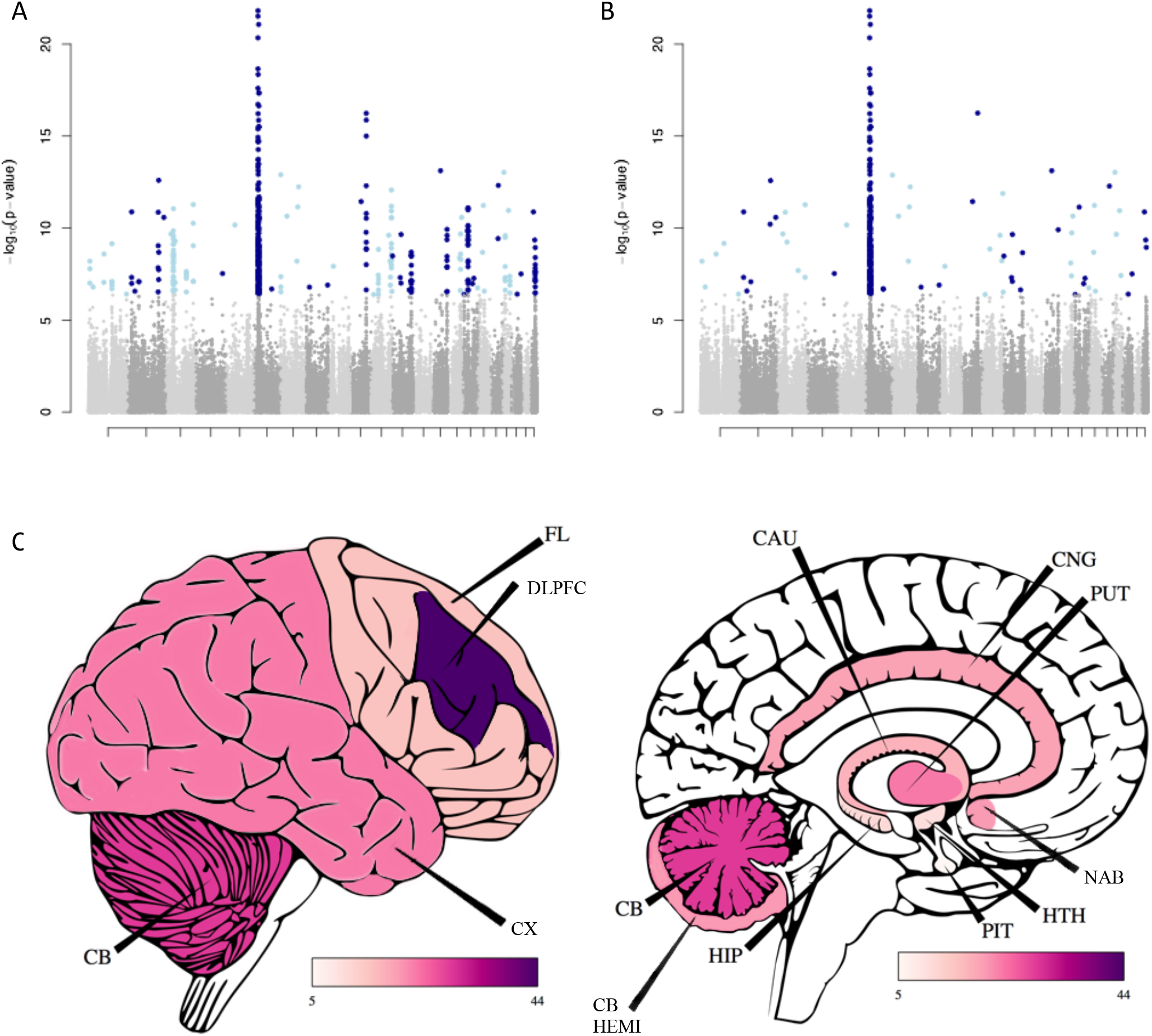
SCZ associations results. A) 413 genes are associated with SCZ across 12 brain tissues B) 67 genes remain significant outside the MHC after stepwise conditional analysis C) Number of genome-wide significant loci, outside the MHC region, identified in each brain region. Abbreviations are as follows; CB-Cerebellum; CX-Cortex; FL-Frontal Cortex; DLPFC-Dorso-lateral pre-frontal cortex; CB HEMI-Cerebellar Hemisphere; HIP-Hippocampus; PIT-Pituitary Gland; HTH-Hypothalamus; NAB-Nucleus Accumbens (Basal Ganglia); PUT-Putamen (Basal Ganglia); CAU-Caudate (Basal Ganglia); CNG-Anterior Cingulate Cortex

To identify the top independent associations within genomic regions, which include multiple associations for a single gene across tissues, or multiple nearby genes, we partitioned genic associations into 58 groups defined based on genomic proximity and applied stepwise forward conditional analysis within each group (Supplementary Table 1). In total, 67 genes remained genome-wide significant after conditioning (Table 1; Figure 3A-B). The largest signal was identified in the CMC DLPFC predicted expression data (24 genes; Figure 3C), followed by the Putamen (7 genes). 19/67 genes did not lie within 1Mb of a previously genome-wide significant GWAS locus^22^ (shown in bold, Table 1); of these, 5/19 genes were within 1Mb of a locus which approached genome-wide significance (p<5×10^−07^). The remaining 14 genes all fall within nominally significant PGC-SCZ GWAS loci (p<8×10^−04^), but did not reach genome-wide significance.

**Table 1:**
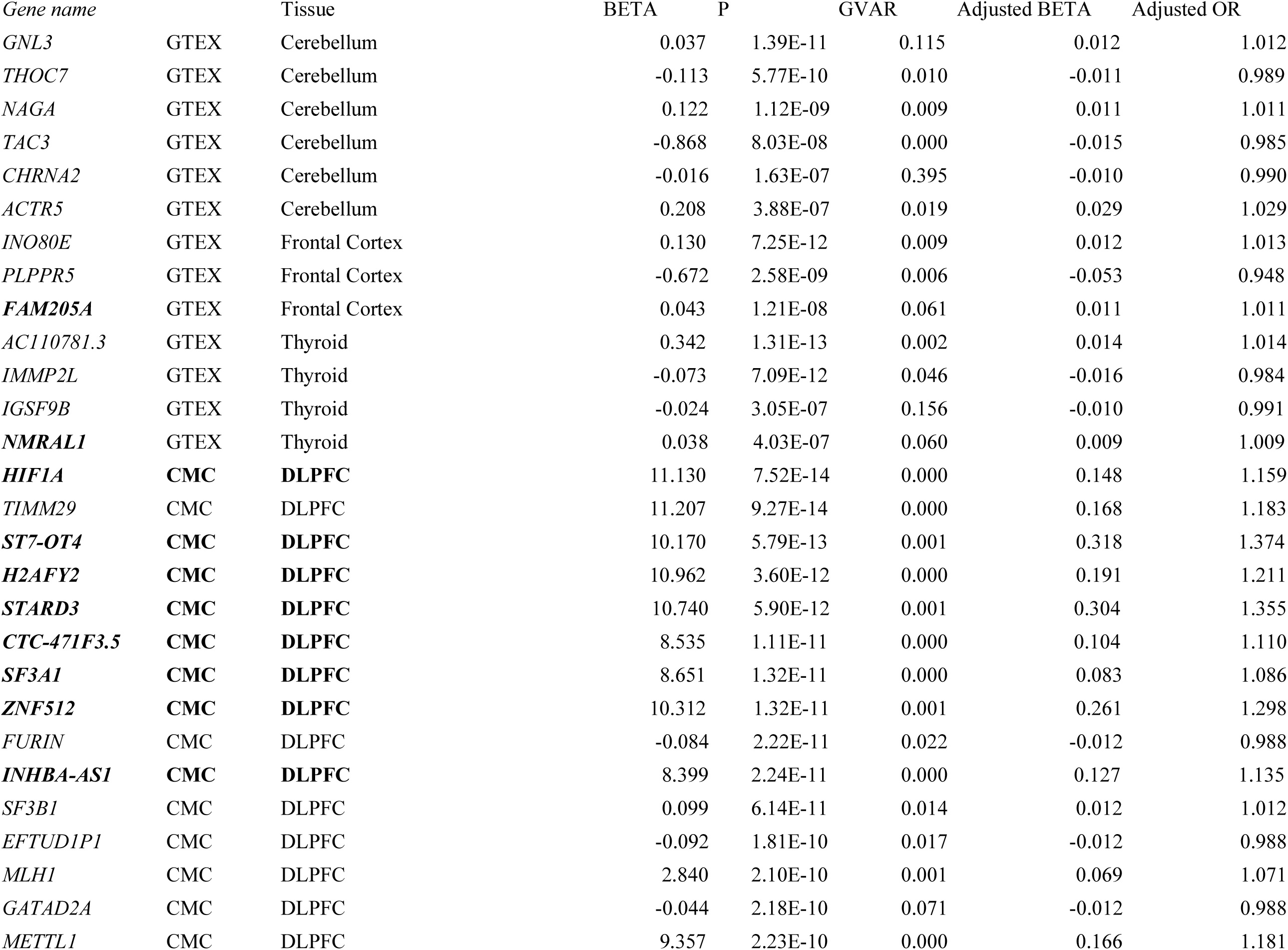
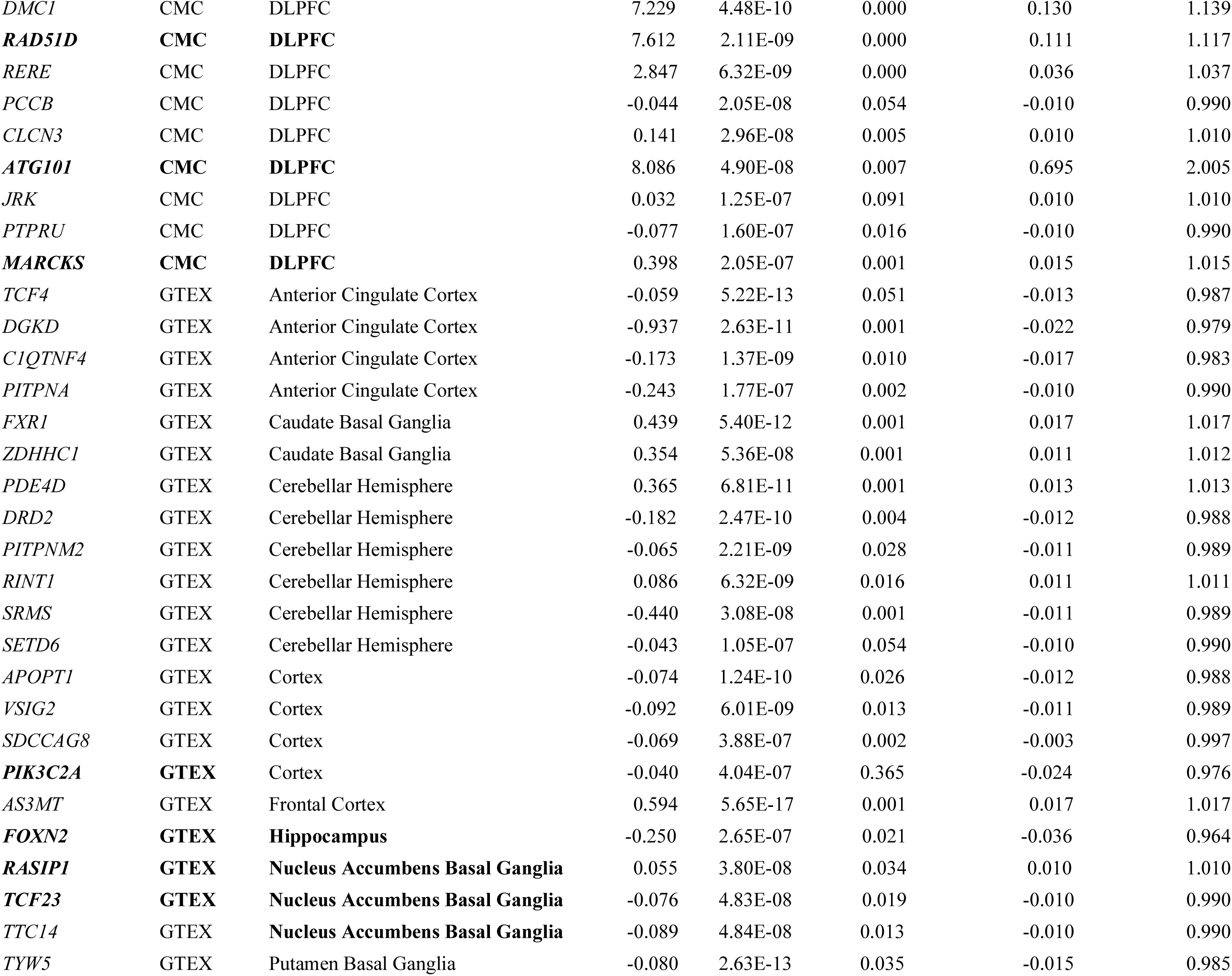
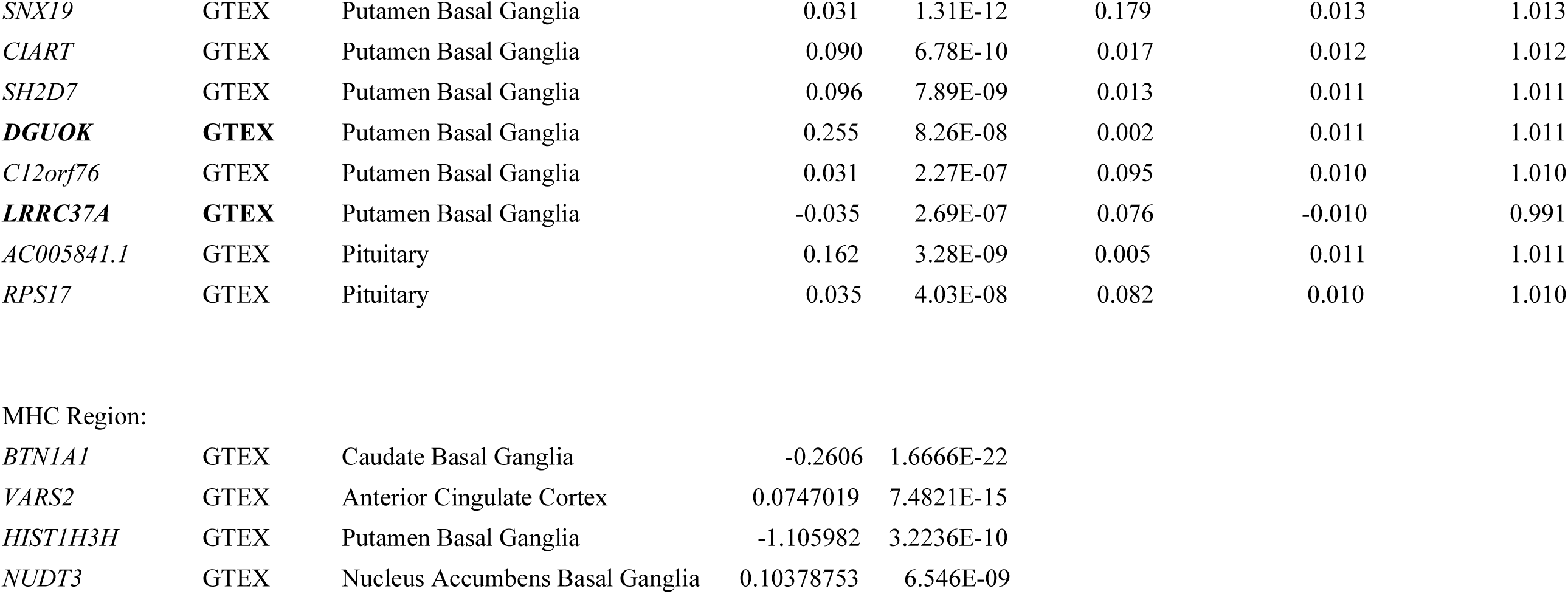
SCZ-associated genes

### Implicated genes highlight SCZ-associated molecular pathways and gene set analyses

We tested for overlap between our non-MHC SCZ-associated genes and 8,657 genesets comprised of 1) hypothesis-driven pathways and 2) general molecular database pathways. We corrected for multiple testing using the Benjamin-Hochberg false discovery rate (FDR) correction^23^.

We identified three significantly associated pathways in our hypothesis-driven analysis (Table 2). Targets of the fragile-X mental retardation protein formed the most enriched pathway (FMRP; p=1.96×10^−8^). Loss of FMRP inhibits synaptic function, is comorbid with autism spectrum disorder, and causes intellectual disability, as well as psychiatric symptoms including anxiety, hyperactivity and social deficits^24^. Enrichment of this large group of genes has been observed frequently, in the original CommonMind analysis^14^, by colleagues investigating the same PGC and CLOZUK2 samples^26^ as well as by investigators studying autism^24,27^. There was a significant enrichment among our SCZ associated genes and genes that have been shown to be intolerant to loss-of-function mutations^28^ (p=5.86×10^−5^) as well as with CNVs associated with bipolar disorder^29^ (p=7.92×10^−8^), in line with a recent variant-based study of the same individuals^26^.

**Table 2:** Significantly enriched pathways and gene sets

| Analysis | Gene Set | Comp P | FDR P |
| --- | --- | --- | --- |
| Hypothesis driven | FMRP-targets | $1.96 \times 10^{-08}$ | $3.097 \times 10^{-06}$ |
| | BP denovo CNV | $7.92 \times 10^{-08}$ | $6.257 \times 10^{-06}$ |
| | HIGH LOF intolerant | $5.86 \times 10^{-05}$ | 0.00309 |
| Agnostic | Increased spleen iron level | $2.72 \times 10^{-08}$ | 0.000245 |
| | Decreased IgM level | $6.80 \times 10^{-07}$ | 0.00307 |
| | Condensed chromosome | $1.99 \times 10^{-06}$ | 0.00598 |
| | Chromosome | $2.80 \times 10^{-06}$ | 0.00632 |
| | Abnormal spleen iron level | $6.79 \times 10^{-06}$ | 0.00765 |
| | Mitotic Anaphase | $6.39 \times 10^{-06}$ | 0.00765 |
| | Mitotic Metaphase and Anaphase | $5.13 \times 10^{-06}$ | 0.00765 |
| | Resolution of Sister Chromatid Cohesion | $5.82 \times 10^{-06}$ | 0.00765 |
| | Increased liver iron level | $1.03 \times 10^{-05}$ | 0.0103 |
| | Separation of Sister Chromatids | $1.28 \times 10^{-05}$ | 0.0115 |
| | Regulation of Rab GTPase activity | $1.78 \times 10^{-05}$ | 0.0123 |
| | Regulation of Rab protein signal transduction | $1.78 \times 10^{-05}$ | 0.0123 |
| | Protein phosphorylated amino acid binding | $1.75 \times 10^{-05}$ | 0.0123 |
| | Chromosome | $2.57 \times 10^{-05}$ | 0.0165 |
| | Hexosaminidase activity | $3.47 \times 10^{-05}$ | 0.0174 |
| | Abnormal learningmemoryconditioning | $3.11 \times 10^{-05}$ | 0.0174 |
| | Abnormal liver iron level | $3.47 \times 10^{-05}$ | 0.0174 |
| | Mitotic Prometaphase | $2.99 \times 10^{-05}$ | 0.0174 |
| | M Phase | $3.70 \times 10^{-05}$ | 0.0176 |
| | Positive regulation of Rab GTPase activity | $5.93 \times 10^{-05}$ | 0.0232 |
| | Rab GTPase activator activity | $5.93 \times 10^{-05}$ | 0.0232 |
| | Protein phosphatase type 2A regulator activity | $5.24 \times 10^{-05}$ | 0.0232 |
| | Replicative senescence | $5.44 \times 10^{-05}$ | 0.0232 |
| | Condensed nuclear chromosome | $7.11 \times 10^{-05}$ | 0.0267 |
|  | Ubiquitin-specific protease activity | 0.000104 | 0.0335 |
| | Ras GTPase activator activity | $9.61 \times 10^{-05}$ | 0.0335 |
|  | Metabolism of porphyrins | 0.000103 | 0.0335 |
|  | Kinetochore | 0.000103 | 0.0335 |
|  | Decreased physiological sensitivity to xenobiotic | 0.000127 | 0.0381 |
|  | Antigen Activates B Cell Receptor Leading to Generation of Second Messengers | 0.000124 | 0.0381 |
|  | Phosphoprotein binding | 0.000146 | 0.0424 |
|  | Abnormal dorsal-ventral axis patterning | 0.000152 | 0.0429 |

Next, we performed an agnostic search for overlap between our schizophrenia-associated genes and ~ 8,500 molecular pathways collated from large, publicly available databases. 33 pathways were significantly enriched after FDR correction (Table 2, Suppl. Table 2), including a number of pathways with some prior literature in psychiatric disease. We identified an enrichment with porphyrin metabolism (p=1.03×10^−4^). Deficiencies in porphyrin metabolism lead to “Porphyria”, an adult-onset metabolic disorder with a host of associated psychiatric symptoms, in particular episodes of violence and psychosis^30–35^. Five pathways potentially related to porphyrin metabolism, regarding abnormal iron level in the spleen, liver and kidney are also significantly enriched, including 2/5 of the most highly enriched pathways (p<2.0 ×10^−04^). The PANTHER and REACTOME pathways for Heme biosynthesis and the GO pathway for protoporphyrinogen IX metabolic process, which are implicated in the development of porphyric disorders, are also highly enriched (p=2.2 ×10^−04^, 2.6 ×10^−04,^ 4.1 ×10^−04^), although do not pass FDR-correction.

Hexosaminidase activity was enriched (p=3.47 ×10^−05^) in our results; this enrichment is not driven by a single highly-associated gene; rather, every single gene in the HEX-A pathway is nominally significant in the SCZ association analysis (Supplementary Table 2). Deficiency of hexosaminidase A (HEX-A) results in serious neurological and mental problems, most commonly presenting in infants as “Tay-Sachs” disease^36^. Adult-onset HEX-A deficiency presents with neurological and psychiatric symptoms, notably including onset of psychosis and schizophrenia^37^. Five pathways corresponding to Ras- and Rab-signaling, protein regulation and GTPase activity were enriched (p<6×10^−05^). These pathways have a crucial role in neuron cell differentiation^38^ and migration^39^, and have been implicated in the development of schizophrenia and autism^40–43^. We also find significant enrichment with protein phosphatase type 2A regulator activity (p=5.24×10^−05^), which was associated with MDD and across MDD, BPD and SCZ in the same large integrative analysis^44^, and has been implicated in antidepressant response and serotonergic neurotransmission^45^.

### Predicted gene expression changes are consistent with functional validation studies

To test the functional impact of our SCZ-associated predicted gene expression changes (GREX), we performed two in-silico analyses. First, we compared directions of effect in our meta-analysis to those in the CMC analysis of differentially expressed genes between SCZ cases and controls. This analysis highlighted six loci where expression levels of a single gene putatively affected schizophrenia risk. All six of these genes are nominally significant in our DLPFC analysis, and two (*CLCN3* and *FURIN*) reach genome-wide significance. In the conditional analysis across all brain regions, one additional gene (*SNX19*) reaches genome-wide significance. The direction of effect for all six genes matches the direction of gene expression changes observed in the original CMC paper, indicating that gene expression estimated in the imputed transcriptome reflects measured expression levels in brains of individuals with Schizophrenia. Further, this observation is consistent with a model where the differential expression signature observed in CMC is caused by genetics rather than environment.

The original CMC analysis identified 21 eSNP genes using SHERLOCK^14,46^, of which 17 were present in our CMC DLPFC analysis. 14/17 genes reached nominal significance (significantly more than expected by chance, p=3.6×10^−16^), and 11 reached genome-wide significance (binomial p-value 6.04×10^−55^). Additionally, 31 regions contained genes ranked highly by Sherlock in the original CMC analysis (supplementary data file 2 in Fromer, M. *et al.* Gene expression elucidates functional impact of polygenic risk for schizophrenia. *Nat. Neurosci.* 19, 1442-1453 (2016)^14^). Of these, 14 regions lay near one of our CMC DLPFC associated genes, and 13/14 regions had common genes between SHERLOCK and PrediXcan analyses. Five loci included multiple SHERLOCK genes; in every instance we are able to specifically identify one or two associated genes from the longer SHERLOCK list.

To understand the impact of altered expression of our 67 SCZ-associated genes, we performed an in-silico analysis of mouse mutants, by collating large, publicly available mouse databases^47–51^. We identified mutant mouse lines lacking expression of 37/67 of our SCZ-associated genes, and obtained 5,333 phenotypic data points relating to these lines, including 1,170 related to behavioral, neurological or craniofacial phenotypes. 25/37 genes were associated with at least one behavioral, neurological or related phenotype (Supplementary table 3). We repeated this analysis for genes identified in 366 GWAS, including any GWAS for which at least ten mutant mouse lines exist (105 GWAS). SCZ-associated genes were more likely to be associated with behavior, brain development and nervous system phenotypes than genes in these GWAS sets (p=0.057).

### Spatiotemporal expression of SCZ-associated genes indicated distinct patterns of risk throughout development

We assessed expression of our SCZ-associated genes throughout development using BRAINSPAN^52^. Data were partitioned into eight developmental stages (four pre-natal, four postnatal), and four brain regions^29,52^ (Figure 4A). We noted that SCZ-associated genes were significantly co-expressed, in both pre-natal and post-natal development and in all four brain regions, based on local connectedness^53^ (Figure 4B), global connectedness^53^ (i.e., average path length between genes, supplementary Figure 6), and network density (i.e., number of edges, supplementary Figure 7). Examining pairwise gene expression correlation (suppl. Fig 8) and gene co-expression networks (suppl. Fig 9) for each spatiotemporal point indicated that the same genes do not drive this co-expression pattern throughout development; rather, it appears that separate groups of genes drive early pre-natal, late pre-natal and post-natal clustering.

**Figure 4:**
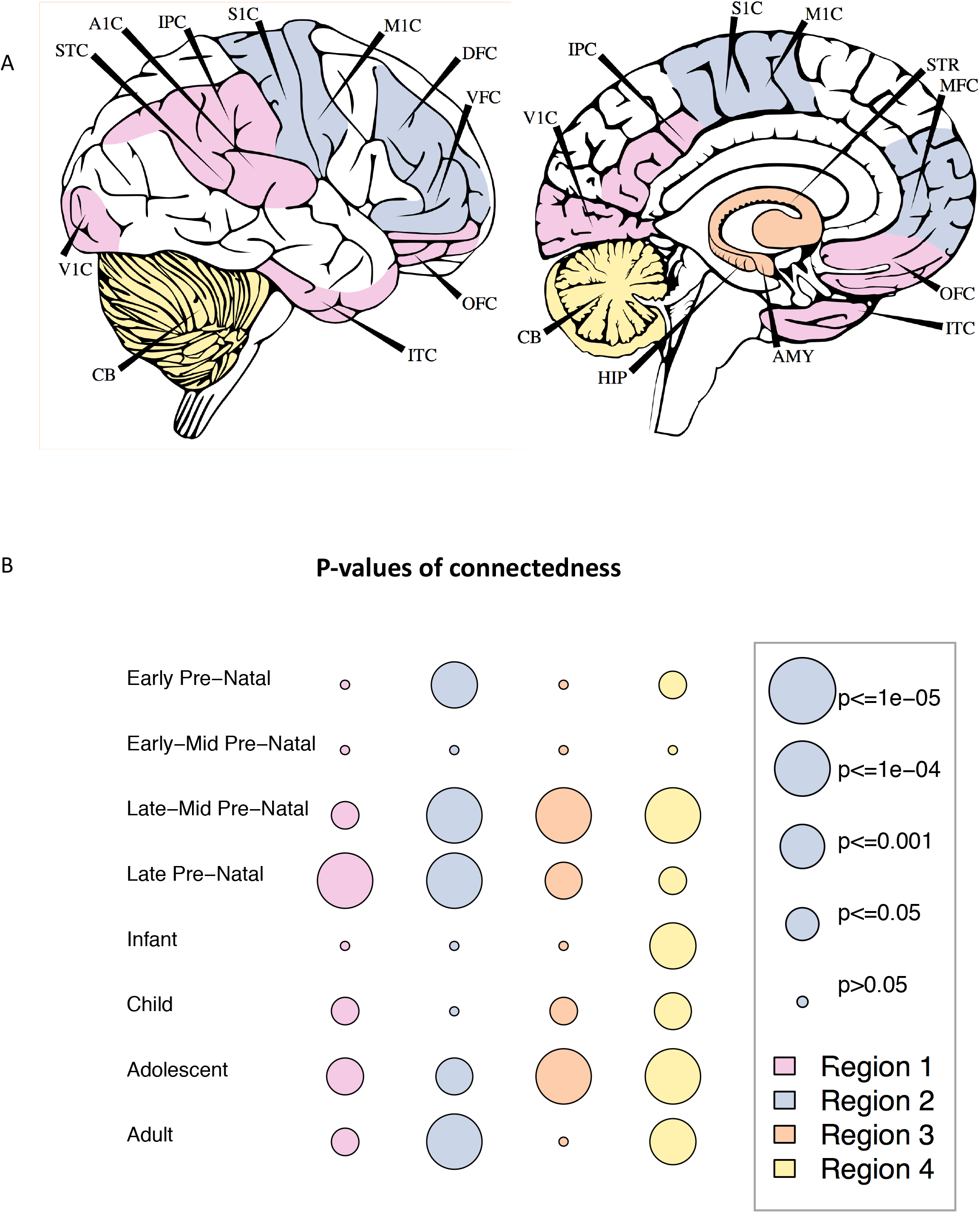
SCZ-associated genes are co-expressed throughout development and across brain regions. A) Brain tissues selected for each of four BRAINSPAN regions. Region 1: IPC, V1C, ITC, OFC, STC, A1C; Region 2:S1C, M1C, DFC, VFC, MFC; Region 3:HIP, AMY, STR; Region 4: CB B) Average clustering coefficients were calculated for all pairs of SCZ-associated genes, and compared to permuted gene networks to obtain empirical significance levels.

To visualize this, we calculated Z scores of gene expression for each SCZ-associated gene, across all 32 time-points (Figure 5). Genes clustered into four groups (supplementary fig 10), with distinct spatio-temporal expression signatures. The largest cluster (Cluster A, Figure 5A; 29 genes) spanned early to late-mid pre-natal development (4-24 weeks post conception), either across the whole brain (22 genes) or in regions 1-3 only (7 genes). 12 genes were expressed in late pre-natal development (Figure 5D; 25-38 pcw); 10 genes were expressed in regions 1-3, post-natally and in the late pre-natal period (Figure 5C), and 15 genes were expressed throughout development (Figure 5B), either specifically in region four (nine genes) or throughout the brain (six genes). We used a stratified qq-plot approach^54^ to examine whether SNPs in cis-regions of genes in these four clusters are differentially enriched in psychiatric disorders. SNPs in cis-regions of genes in the two pre-natal clusters are more highly enriched than SNPs in cis-regions of genes in post-natal clusters, and compared to all SNPs, in childhood-onset disorders (ASD and ADHD, supplementary figure 13), but not adult-onset disorders (BPD and MDD, data not shown).

**Figure 5:**
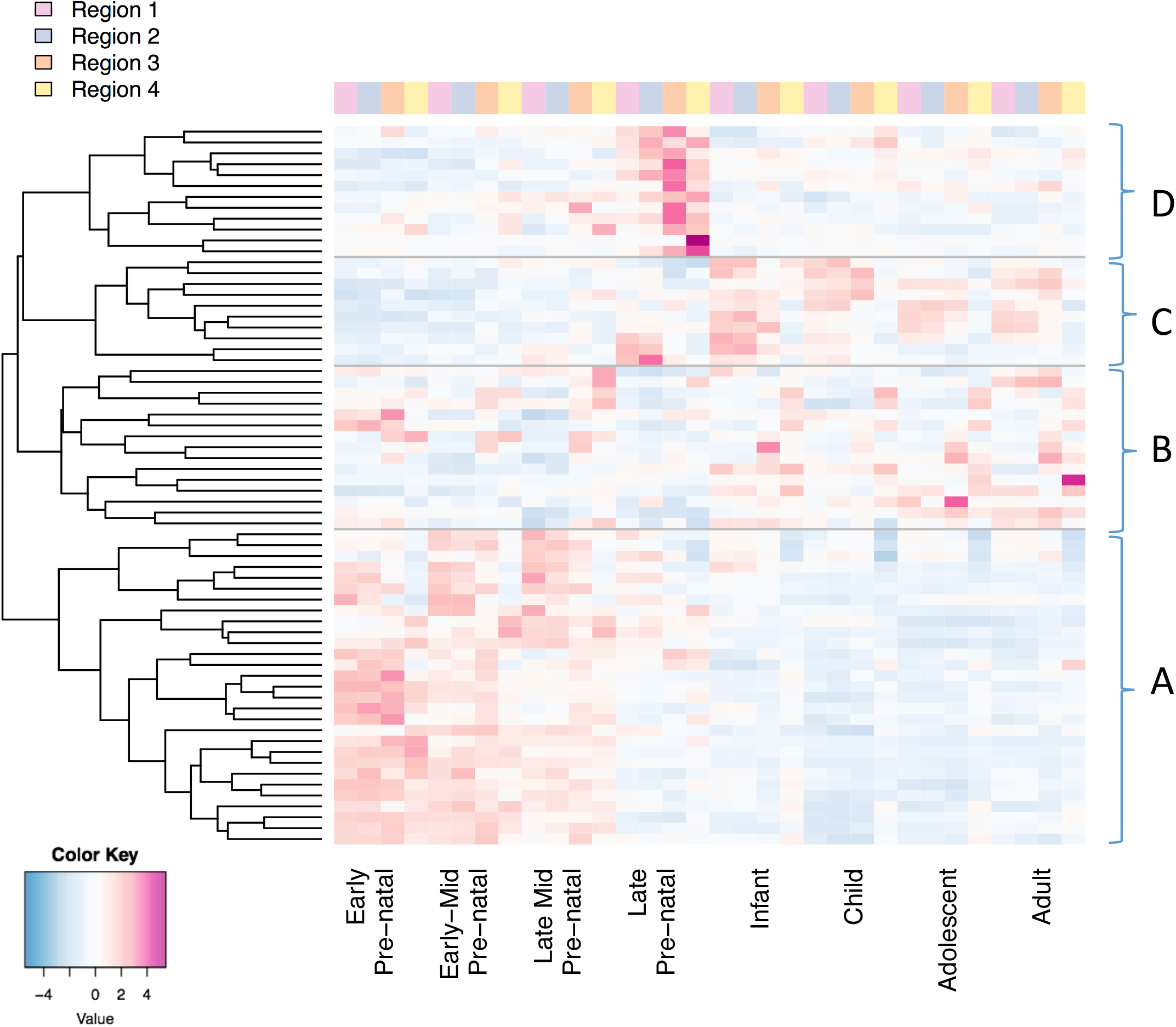
Gene expression patterns for SCZ-associated genes cluster into four groups, relating to distinct spatiotemporal expression. Brain regions are shown in figure 5a. A. 29 genes are expressed in the early-mid pre-natal period (4-24 post-conception weeks) B. 15 genes are expressed throughout development; sub-clusters correspond to either specific expression in region 4, or expression across the brain C. Ten genes are expressed in the late-prenatal (25-38pcw) and post-natal period D. 12 genes are expressed in the late pre-natal period (25-39pcw)

We noticed a relationship between patterns of gene expression and the likelihood of behavioral, neurological or related phenotypes in our mutant mouse model database. Mutant mice lacking genes expressed exclusively pre-natally in humans, or genes expressed pre- and post-natally, were more likely to have any behavioral or neurological phenotypes than mutant mice lacking expression of genes expressed primarily in the third trimester or post-natally (p=1.7×10^−04^) (supplementary figure 11).

## Discussion

In this study, we present gene expression prediction models for the dorso-lateral pre-frontal cortex (DLPFC), constructed using CommonMind Consortium genotype and gene expression data. These prediction models may be applied to either raw data or summary statistics, in order to yield gene expression information in large data sets, and across a range of tissues. This has the significant advantage of allowing researchers to access transcriptome data for non-peripheral tissues, at scales currently prohibited by the high cost of RNA sequencing, and circumventing distortions in measures of gene expression stemming from errors of measurement or environmental influences. Since disease status may alter gene expression but not the germline profile, analyzing genetically regulated expression ensures that we identify only the causal direction of effect between gene expression and disease^15^. Large, imputed transcriptomic datasets represent the first opportunity to study the role of subtle gene expression changes (and therefore modest effect sizes) in disease development.

There are some inherent limitations to this approach. The accuracy of transcriptomic imputation (TI) is reliant on access to large eQTL reference panels, and it is therefore vital that efforts to collect and analyze these samples continue. TI has exciting advantages for gene discovery as well as downstream applications^15,55,56^; however, the relative merits of existing methodologies are as yet under-explored. Our analysis suggests that, overall, sparser elastic net models better capture gene expression regulation than BSLMM; at the same time, the improved performance of elastic net over max-eQTL models suggests that a single eQTL model is over-simplified^2,15^. Fundamentally, transcriptomic imputation methods model only the genetically regulated portion of gene expression, and so cannot capture or interpret variance of expression induced by environment or lifestyle factors, which may be of particular importance in psychiatric disorders. Given the right study design, analyzing genetic components of expression together with observed expression could open doors to better study the role of gene expression in disease.

Sample size and tissue matching contribute to accuracy of TI results. Our CMC-derived DLPFC prediction models had higher average validation R^2^ values in external DLPFC data than GTEx-derived brain tissue models. Notably, the model with the second highest percent of genes passing the R^2^ threshold is the Thyroid, which has the largest sample size among the GTEx brain prediction models. When looking at mean R^2^ values, the second highest value comes from the GTEx Frontal Cortex, despite the associated small sample size, implying at least some degree of tissue specificity of eQTLs architecture.

We were able to compare TI accuracy in European and African-American individuals, and found that our models were applicable to either ethnicity with only a small decrease in accuracy. Common SNPs shared across ethnicities have important effects on gene expression, and as such we expect GREX to have consistency across populations. There is a well-documented dearth of exploration of genetic associations in non-European cohorts^57,58^ We believe that these analyses should be carried out in non-European cohorts.

We applied the CMC DLPFC prediction models, along with 12 GTEx-derived brain expression prediction models, to schizophrenia cases and controls from the PGC2 and CLOZUK2 collections, constituting the largest transcriptomic analysis of schizophrenia to date. Predicted gene expression levels were calculated for 19,661 unique genes across brain regions (Figure 1C) and tested for association with SCZ case-control status. We identified 413 significant associations, constituting 67 independent associations. We found significant replication of our CMC DLPFC associations in a large independent replication cohort, in collaboration with the iPSYCH-GEMS consortium. A recent TWAS study of 30 GWAS summary statistic traits^55^ identified 38 non-MHC genes associated at tissue-level significance with SCZ in CMC- and GTEx-derived brain tissues (ie, matching those used in our study). Of these, 26 also reach genome-wide significance in our study, although in many instances these genes are not identified as the lead independent associated gene following our conditional analysis. Among our 67 SCZ-associated genes, 19 were novel, i.e. did not fall within 1Mb of a previous GWAS locus (including 5/7 of the novel brain genes identified in the recent TWAS analysis).

We used conditional analyses to identify independent associations within loci. These analyses clarify the most strongly associated genes and tissues (Table 1), while we note that nearly collinear gene-tissue pairs could also represent causal associations. The tissues highlighted allowed us to tabulate apparently independent contributions to SCZ risk from different brain regions, even though their transcriptomes are highly correlated generally. We find DLPFC and Cerebellum effects, as well as from Putamen, Caudate and Nucleus Accumbens Basal Ganglia.

We used these genic associations to search for enrichments with molecular pathways and gene sets, and identified 36 significant enriched pathways. Among novel pathways, we identified a significant association with HEX-A deficiency. Despite the well-studied and documented symptomatic overlap between adult-onset HEX-A deficiency and schizophrenia, we believe that this is the first demonstration of shared genetics between the disorders. Notably, this overlap is not driven by a single highly-associated gene which is shared by both disorders; rather, every single gene in the HEX-A pathway is nominally significant in the SCZ association analysis, and five genes have p < 1×10^−03^, indicating that there may be substantial shared genetic aetiology between the two disorders that warrants further investigation. Additionally, we identified a significant overlap between our SCZ-associated genes and a number of pathways associated with porphyrin metabolism. Porphyric disorders have been well characterized and are among early descriptions of “schizophrenic” and psychotic presentations of schizophrenia, as described in the likely eponymous mid-19^th^ century poem “Porphyria’s Lover”, by Robert Browning^59^, and have been cited as a likely diagnosis for the various psychiatric and metabolic ailments of Vincent van Gogh^60–65^ and King George III^66^.

Finally, we assessed patterns of expression for the 67 SCZ-associated genes throughout development using spatio-temporal transcriptomic data obtained from BRAINSPAN. We identified four clusters of genes, with expression in four distinct spatiotemporal regions, ranging from early pre-natal to strictly post-natal expression. There are plausible hypotheses and genetic evidence for SCZ disease development in adolescence, given the correlation with age of onset, as well as prenatally, supported by genetic overlap with neurodevelopmental disorders^67–69^ as well as the earlier onset of cognitive impairments^70–73^. Understanding the temporal expression patterns of SCZ-associated genes can help to elucidate gene development and trajectory, and inform research and analysis design. Identification of SCZ-associated genes primarily expressed prenatally is striking given our adult eQTL reference panels, and may reflect common eQTL architecture across development, which is known to be partial^74–76^; therefore, our results should spur interest in extending TI data and/or methods to early development^74^. Identification of SCZ-associated genes primarily expressed in adolescence and adult-hood is of particular interest for direct analysis of the brain transcriptome in adult psychiatric cases.

eQTL data have been recognized for nearly a decade as potentially important for understanding complex genetic variation. Nicolae et al^1^ showed that common variant-common disease associations are strongly enriched for genetic regulation of gene expression. Therefore, integrative approaches combining transcriptomic and genetic association data have great potential. Current TI association analyses increase power for genetic discovery, even while many open areas of TI remain to be developed, such as leveraging additional data types such as chromatin modifications^77^ (e.g. methylation, histone modification), imputing different tissues or different exposures (e.g. age, smoking, trauma) and modeling trans/coexpression effects. It remains critical to leverage TI associations to provide insights into specific disease mechanisms. Here, the accelerated identification of disease associated genes allows the detection of novel pathways and distinct spatiotemporal patterns of expression in schizophrenia risk.

## Online Methods (Limit 3,000 words, at end of manuscript, currently 2,064)

### Creating gene expression predictors for the dorso-lateral pre-frontal cortex eQTL Data

Genotype and RNAseq data were obtained for 538 European individuals through the CommonMind Project^14^. RNA-seq data were generated from post-mortem human dorsolateral prefrontal cortex (DLPFC). The gene expression matrix was normalized to log(counts per million) using voom. Adjustments were made for known covariates (including sample ascertainment, quality, experimental parameters, ancestry) and surrogate variables, using linear modelling with voom-derived regression weights. Details on genotyping, imputation and RNA-seq generation may be found in the CommonMind Consortium flagship paper^14^.

A 1% MAF cut-off was applied. Variants were filtered to remove any SNPs in high LD (R^2^>0.9), indels, and all variants with ambiguous ref/alt alleles. All protein coding genes on chromosomes 1-22 with at least one cis-SNP after these QC steps were included in this analysis. SNPs in trans have been shown not to provide a substantial improvement in prediction accuracy^15^ and were not included here.

### Building gene expression prediction databases

Gene expression prediction models were created following the “PrediXcan” method^15^. Matched genotype and gene expression data were used to identify a set of variants that influence gene expression (Supplementary Figure 2A). Weights for these variants are calculated using regression in a ten-fold cross-validation framework. All cross-validation folds were balanced for diagnoses, ethnicity, and other clinical variables.

All SNPs within the cis-region (+/-1mb) of each gene were included in the regression analysis. Accuracy of prediction was estimated by comparing predicted expression to measured expression, across all 10 cross-validation folds; this correlation was termed cross-validation R^2^ or R_CV_^2^. Genes with R_CV_^2^ > 0.01 (~p<0.05) were included in our final predictor database.

Prediction models were compared across four different regression methods; elastic net (prediXcan), ridge regression (using the TWAS method^16^), Bayesian sparse linear mixed modelling (BSLMM; TWAS), and linear regression using the best eQTL for each gene (Supplementary Figure 1A). Mean R_CV_^2^ values were significantly higher for elastic net regression (mean R_CV_^2^=0.056) than for eQTL-based prediction (mean R_CV_^2^=0.025), BSLMM (mean R_CV_^2^=0.021) or Ridge Regression (mean R_CV_^2^=0.020). The distribution of R_CV_^2^ values was also significantly higher for elastic net regression than for any other method (ks-test, p<2.2×10^−16^).

### Replication of gene expression prediction models in independent data

Predictive accuracy of CMC DLPFC models were tested in two independent datasets. First, we used data from the Religious Orders Study and Memory and Aging Project (ROSMAP^19^). This study included genotype data and DLPFC RNA-seq data^78^ for 451 individuals of European descent (Supplementary Figure 2B).

DLPFC genetically-regulated expression (GREX) was calculated using the CMC DLPFC predictor models. Correlation between RNA-seq expression and CMC DLPFC GREX (“Replication R^2^ values” or R_R_^2^) was used as a measure of predictive accuracy. R_R_^2^ was calculated including correction for ten ancestry components, as follows:

*Equation 1: R_R_^2^ calculation.*

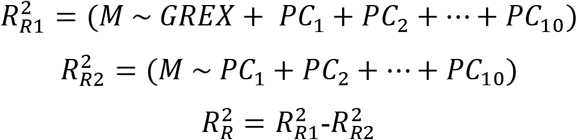

Where:

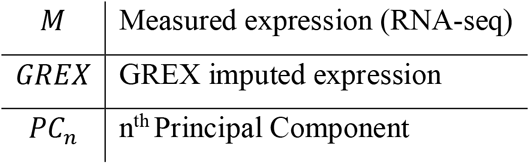

A small number of genes (158) had very low predictive accuracy and were removed from further analyses. Cross-validation R^2^ (R_CV_^2^) values and R_R_^2^ values were highly correlated (rho=0.62, p<2.2e-16; Supplementary Figure 3A). 55.7% of CMC DLPFC genes had R_R_^2^ values > 0.01.

Prediction accuracy was also assessed for 11 publicly available GTEx neurological predictor databases, and R_R_^2^ values used to compare to CMC DLPFC performance. CMC DLPFC models had higher average R_R_^2^ values, more genes with R_R_^2^ > 0.01, and significantly higher overall distributions of R_R_^2^values than any of the twelve GTEx brain tissue models (ks-test, p<2.2e-16; Figure 1A,B).

To estimate trans-ancestral prediction accuracy, genetically regulated gene expression was calculated for 162 African-American individuals and 280 European individuals from the NIMH Human Brain Collection Core (HBCC) dataset^79^ (Supplementary Figure 2C). Predicted gene expression levels were compared to DLPFC expression levels measured using microarray. There was a significant correlation between the European and African-American samples for R_CV_^2^ values and R_R_^2^ values (rho=0.66, 0.56; Supplementary figure 3B-C). R_R_^2^ values were higher on average in Europeans, but were significantly correlated between African-Americans and Europeans (rho=0.78, p<2.2e-16, Pearson test; supplementary figure 3D).

### Extension to Summary Statistics

Transcriptomic Imputation may be applied to summary statistics instead of raw dosages, in instances where raw data is unavailable. However, this method suffers from slightly reduced accuracy, requires covariance matrices calculated in an ancestrally-matched reference population^80^ (usually only possible for European cohorts), and precludes testing of endophenotypes within the data, and so should not be applied when raw data is available.

We assessed concordance between CMC DLPFC transcriptomic imputation results using summary-statistics (MetaXcan^80^) and raw genotypes (PrediXcan^15^) using nine European and three Asian PGC-SCZ cohorts^22^ for which both data types were available. Cohorts were chosen to encompass a range of case: control ratios, to test previous suggestions that accuracy is reduced in unbalanced cohorts^80^. Covariances for all variants included in the DLPFC predictor models were computed using MetaXcan^80^. For all European cohorts, Pearson correlation of log-10 p-values and effect sizes was above 0.95. The mean correlation was 0.963 (Supplementary Figure 4). There was no correlation between total sample size, case-control ratio, p-value or effect-size. Seven genes were removed due to discordant p-values. For the three Asian cohorts tested, the mean correlation was 0.91 (Supplementary Figure 5).

Concordance was also tested for the same nine European PGC-SCZ cohorts, across 12 neurological GTEx prediction databases. All correlations were significant (rho>0.95, p<2.2e-16). There was a significant correlation between p-value concordance and case-control ratio (rho=0.37, p=7.606 ×10^−15^). 114 genes had discordant p-values between the two methods and were excluded from future analyses.

## Application to Schizophrenia

### Dataset Collection

We obtained 53 discovery cohorts for this study, including 40,299 SCZ cases and 65,264 controls (Figure 2). 52/53 cohorts (35,079 cases, 46,441 controls) were obtained through collaboration with the Psychiatric Genomics Consortium, and are described in the 2014 PGC Schizophrenia GWAS^22^. The remaining cohort, referred to as CLOZUK2, constitutes the largest single cohort of individuals with Schizophrenia (5,220 cases and 18,823 controls), collected as part of an effort to investigate treatment-resistant Schizophrenia^26^.

50/53 datasets included individuals of European ancestry, while three datasets include individuals of Asian ancestry (1,836 cases, 3,383 controls). All individuals were ancestrally matched to controls. Information on genotyping, quality control and other data management issues may be found in the original papers describing these collections^22,26^. All sample collections complied with ethical regulations. Details regarding ethical compliance and consent procedures may be found in the original manuscripts describing these collections^22,26^.

Access to dosage data was available for 44/52 PGC-SCZ cohorts. The remaining PGC cohorts, and the CLOZUK2 cohort provided summary statistics. Three European PGC cohorts were trio-based, rather than case-control.

Additionally, we tested for replication of our CMC DLPFC associations in an independent dataset of 4,133 cases and 24,788 controls obtained through collaboration with the iPSYCH-GEMS schizophrenia working group (effective sample size 14,169.5; Figure 2B, supplementary information).

### Transcriptomic Imputation and association testing

Transcriptomic Imputation was carried out individually for each case-control PGC-SCZ cohort with available dosage data (44/52 cohorts). Predicted gene expression levels were computed using the DLPFC predictors described in this manuscript, as well as for 11 other brain tissues prediction databases created using GTEx tissues^15,20,21,81^ (Figure 1C). Associations between predicted gene expression values and case-control status were calculated using a linear regression test in R. Ten ancestry principal components were included as covariates. Association tests were carried out independently for each cohort, across 12 brain tissues.

For the 8 PGC cohorts with no available dosage data, the three PGC trio-based analyses, and the CLOZUK2 cohort, a summary-statistic based transcriptomic imputation approach was used (“MetaXcan”), as described previously.

### Meta-analysis

Meta-analysis was carried out across all 53 cohorts using METAL^82^. Cochran’s Q test for heterogeneity was implemented in METAL^82,83^, and a heterogeneity p-value threshold of p > 1×10^−3^ applied to results. A conservative significance threshold was applied to these data, correcting for the total number of genes tested across all tissues (121,611 gene-region tests in total). This resulted in a genome-wide significance threshold of 4.1×10^−7^.

Effect sizes and direction of effect quoted in this manuscript refer to changes in predicted expression in cases compared to controls i.e., genes with negative effect sizes have decreased predicted expression in cases compared to controls.

### Identifying independent associations

We identified a number of genomic regions which contained multiple gene associations and/or genes associated across multiple tissues. We identified 58 of these regions, excluding the MHC, based on distance between associated genes, and verified them using visual inspection. In order to identify independent genic associations within these regions, we carried out a stepwise forward conditional analysis following “GCTA-COJO” theory^84^ using “CoCo” (https://github.com/theboocock/coco/), an R implementation of GCTA-COJO. CoCo allows the specification of custom correlation matrices by the user (for example, ancestrally specific LD matrices). For each region, we generated a predicted gene expression correlation matrix for all significant genes (p≤1×10^−6^), as the root-effective sample size (Neff, eqn 2) weighted average correlation across all cohorts where we had access to dosage data.

*Equation 2: Effective Sample Size, N_eff_*

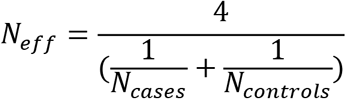

Forward stepwise conditional analysis of all significant genes was carried out using joint linear regression modeling. First, the top-ranked gene was added to the model, then the next most significant gene in a joint model is added if significant at a given p-value threshold, and so on until either all genes are added to the model, or no joint statistic reaches the significance threshold.

We calculated effect sizes and odds ratios for SCZ-associated genes by adjusting “CoCo” betas to have unit variance (Table 1, eqn. 3).

*Equation 3: GREX Beta adjustment*

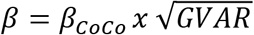

Where GVAR is the variance of the GREX predictor for each gene.

### Gene set Analyses

Pathway analyses were carried out using an extension to MAGMA^85^. P-values were assigned to genes using the most significant p-value achieved by each gene in the meta-analysis. We then carried out a competitive gene-set analysis test using these p-values, using two gene sets:

1. 159 gene sets with prior hypotheses for involvement in SCZ development, including loss-of-function intolerant genes, CNV-intolerant genes, targets of the fragile-X mental retardation protein, CNS related gene sets, and 104 behavioural and neurological pathways from the Mouse Genome Informatics database^14,26,67,86^.
2. An agnostic analysis, including ~8,500 gene sets collated from publicly available databases including GO^87,88^, KEGG^89^, REACTOME^90^, PANTHER^91,92^, BIOCARTA^93^ and MGI^48^. Sets were filtered to include only gene sets with at least ten genes.

Significance levels were adjusted across all pathways included in either test using the Benjamini-Hochberg “FDR” correction in R^23^.

### Coexpression of SCZ genes throughout development

We investigate spatiotemporal expression of our associated genes using publicly available developmental transcriptome data, obtained from the BRAINSPAN consortium^94^. We partitioned these data into biologically relevant spatio-temporal data sets^95^, corresponding to four general brain regions; the frontal cortex, temporal and parietal regions, sensory-motor regions, and subcortical regions (Figure 4A^96^), and eight developmental time-points (four pre-natal, four post-natal)^95^.

First, we tested for correlation of gene expression for all SCZ-associated genes at each spatiotemporal time-point. Genes with pearson correlation coefficients >= 0.8 or <=−0.8 were considered co-expressed. 100,000 iterations of this analysis were carried out using random gene sets with equivalent expression level distributions to the SCZ-associated genes. For each gene set, a gene co-expression network was created, with edges connecting all co-expressed genes. Networks were assessed using three criteria; first, the number of edges within the network, as a crude measured of connectedness; second, the Watts-Strogatz average path length between nodes, as a global measure of connectedness across all genes in the network^53^; third, the Watts-Strogatz clustering coefficient, to measure tightness of the clusters within the network^53^. For each spatio-temporal time point, we plotted gene-pair expression correlation (suppl. Fig 8) and co-expression networks (suppl. Fig 9).

For each of the 67 SCZ-associated genes, we calculated average expression at each spatiotemporal point. We then calculated Z-Score of expression specificity using these values, and plotted Z-Scores to visually examine patterns of gene expression throughout development and across brain regions. Clusters were formally identified using a dendrogram cut at height 10 (Suppl. Fig 10).

### In-silico replication of SCZ-associated genes in mouse models

We downloaded genotype, knock-out allele information and phenotyping data for ~10,000 mouse mutant models from five large mouse phenotyping and genotyping projects; Mouse Genome Informatics (MGI^48^), EuroPhenome^47,97^, Mouse Genome Project (MGP^47,49^), International Mouse Phenotyping Consortium (IMPC^50^), and Infection and Immunity Immunophenotyping (3I^98^). Where possible, we also downloaded raw phenotyping data regarding specific assays. In total, we obtained 175,012 phenotypic measurements, across 10,288 mutant mouse models. We searched for any mouse lines with phenotypes related to behavior (natural, observed, stereotypic or assay-induced); cognition or working memory; brain, head or craniofacial dysmorphology; retinal or eye morphology, and/or vision or visual dysfunction or impairment; ear morphology or hearing dysfunction or impairment; neural tube defects; brain and/or nervous system development; abnormal nociception.

We compared the prevalence of psychiatric phenotypes in mutant mice for our SCZ-associated genes to the prevalence among other disease-associated gene sets. We selected 366 GWAS gene sets, and removed any for which fewer than ten mutant mouse models were included in our databases, leaving 105 gene sets. We compared the prevalence of 13 different categories of psychiatric phenotypes, relating to adrenal gland, behavior, brain development, craniofacial dysmorphology, ear/auditory phenotypes, eye dysmorphology, head dysmorphology, nervous system development, abnormal nociception, seizures, thyroid gland, vision phenotypes. For each GWAS gene set, we counted the number of categories with at least one phenotype, and compared to the number in our SCZ-associated gene set to obtain an empirical p-value.

## Data Availability

Our CMC-derived DLPFC prediction models will be made publicly available.

## Acknowledgements

Data were generated as part of the CommonMind Consortium supported by funding from Takeda Pharmaceuticals Company Limited, F. Hoffman-La Roche Ltd and NIH grants R01MH085542, R01MH093725, P50MH066392, P50MH080405, R01MH097276, RO1-MH-075916, P50M096891, P50MH084053S1, R37MH057881 and R37MH057881S1, HHSN271201300031C, AG02219, AG05138 and MH06692.

Brain tissue for the study was obtained from the following brain bank collections: the Mount Sinai NIH Brain and Tissue Repository, the University of Pennsylvania Alzheimer’s Disease Core Center, the University of Pittsburgh NeuroBioBank and Brain and Tissue Repositories and the NIMH Human Brain Collection Core. CMC Leadership: Pamela Sklar, Joseph Buxbaum (Icahn School of Medicine at Mount Sinai), Bernie Devlin, David Lewis (University of Pittsburgh), Raquel Gur, Chang-Gyu Hahn (University of Pennsylvania), Keisuke Hirai, Hiroyoshi Toyoshiba (Takeda Pharmaceuticals Company Limited), Enrico Domenici, Laurent Essioux (F. Hoffman-La Roche Ltd), Lara Mangravite, Mette Peters (Sage Bionetworks), Thomas Lehner, Barbara Lipska (NIMH).

ROSMAP study data were provided by the Rush Alzheimer’s Disease Center, Rush University Medical Center, Chicago. Data collection was supported through funding by NIA grants P30AG10161, R01AG15819, R01AG17917, R01AG30146, R01AG36836, U01AG32984, U01AG46152, the Illinois Department of Public Health, and the Translational Genomics Research Institute.

The iPSYCH-GEMS team would like to acknowledge funding from the Lundbeck Foundation (grant no R102-A9118 and R155-2014-1724), the Stanley Medical Research Institute, an Advanced Grant from the European Research Council (project no: 294838), the Danish Strategic Research Council the Novo Nordisk Foundation for supporting the Danish National Biobank resource, and grants from Aarhus and Copenhagen Universities and University Hospitals, including support to the iSEQ Center, the GenomeDK HPC facility, and the CIRRAU Center.

The Genotype-Tissue Expression (GTEx) Project was supported by the Common Fund of the Office of the Director of the National Institutes of Health, and by NCI, NHGRI, NHLBI, NIDA, NIMH, and NINDS. The data used for the analyses described in this manuscript were obtained from the GTEx Portal on 09/05/16. BrainSpan: Atlas of the Developing Human Brain [Internet]. Funded by ARRA Awards 1RC2MH089921-01, 1RC2MH090047-01, and 1RC2MH089929-01.

